# Expression of lamina proteins Lamin Dm0 and Kugelkern suppresses stem cell proliferation

**DOI:** 10.1101/177998

**Authors:** Roman Petrovsky, Jörg Großhans

**Author notes:** Summary statement: Expression of Lamin Dm0 or Kugelkern curb stem cell proliferation by suppression of Jak/Stat signalling.

## Abstract

The nuclear lamina is involved in numerous cellular functions, such as gene expression, nuclear organization, nuclear stability, and cell proliferation. The mechanism underlying the involvement of lamina is often not clear, especially in physiological contexts. Here we investigate how the farnesylated lamina proteins Lamin Dm0 and Kugelkern are linked to proliferation control of intestinal stem cells (ISCs) in adult *Drosophila* flies by loss-of-function and gain-of-function experiments. We found that ISCs mutant for Lamin Dm0 or Kugelkern proliferate, whereas overexpression of Lamin Dm0 or Kugelkern strongly suppressed proliferation. The anti-proliferative activity is, at least in part, due to suppression of Jak/Stat but not Delta/Notch signalling. Lamin Dm0 expression suppresses Jak/Stat signalling by normalization of about 50% of the Stat target genes in ISCs.

**Author summary:** The nuclear lamina is a protein meshwork that lies beneath the inner side of the nuclear membrane and interacts with nuclear pores, chromatin and the cytoskeleton. Changes in proteins of the nuclear lamina cause a wide range of diseases which are often not well understood. It is hypothesized that impairment of stem cell function, as a result of lamina changes, might play a key role in some of those diseases. Here we use the well characterized *Drosophila* midgut as a system to investigate the role of lamina proteins Lamin Dm0 and Kugelkern on stem cell proliferation.

## 1 Introduction

In *Drosophila*, induced expression of farnesylated Lamin Dm0 (B type lamin) and Kugelkern (Kuk) promote nuclear deformations, DNA damage, reduced heterochromatin and reduced lifespan [4, 1]. While Lamin Dm0 is a typical member of the lamin family Kuk is structurally distinct from lamins and intermediate filament proteins. However similar to lamins it contains a putative coiled coil motif in its N-terminus, a nuclear localization signal (NLS) and a CaaX motif in the C-terminus that facilitates its farnesylation [5]. Functionally Kuk is related to lamins, since expression of Kuk induces similar phenotypes as expression of farnesylated lamins in several cellular systems [5, 4, 28], including abnormal nuclear morphology, decreased heterochromatin and increased DNA damage [5, 4, 28].

Expression of farnesylated lamina proteins prelamin A and Progerin has been shown to effect the proliferation and number of stem cells in mice [12, 33]. We wondered if similar effects are found in *Drosophila* upon expression of Lamin Dm0 and Kuk and what molecular mechanisms might facilitate these effects.

Adult *Drosophila* flies contain three proliferative populations of stem cells, stem cells in the germ line, associated somatic follicle cells and intestinal stem cells in the midgut epithelium (ISC) [38, 36, 17, 23, 24]. The midgut has a simple morphological organization and only five major cell types: 1. Intestinal stem cells (ISCs), precursors of EEs and EBs and the main proliferating cells in the midgut [18, 15] 2. Enteroblasts (EBs), progenitor cells for ECs, 3. absorptive enterocytes (ECs) 4. secretory enteroendocrine cells (EEs), 5. viceral muscle cells (Fig. 1A). These five cell types can be genetically and histologically distinguished [21, 25, 23, 24]. Additional cell types are found in the copper cell region of the midgut which is not topic of this work. The level of proliferation during homeostasis (standard conditions in the laboratory) is a few mitotic events per gut and a full turnover of the tissue within about two weeks, in females [16]. Following infection with bacteria, the gut regenerates involving a strong induction of ISC proliferation (up to 10 times higher) and rapid tissue turnover [7].

**Figure 1.**
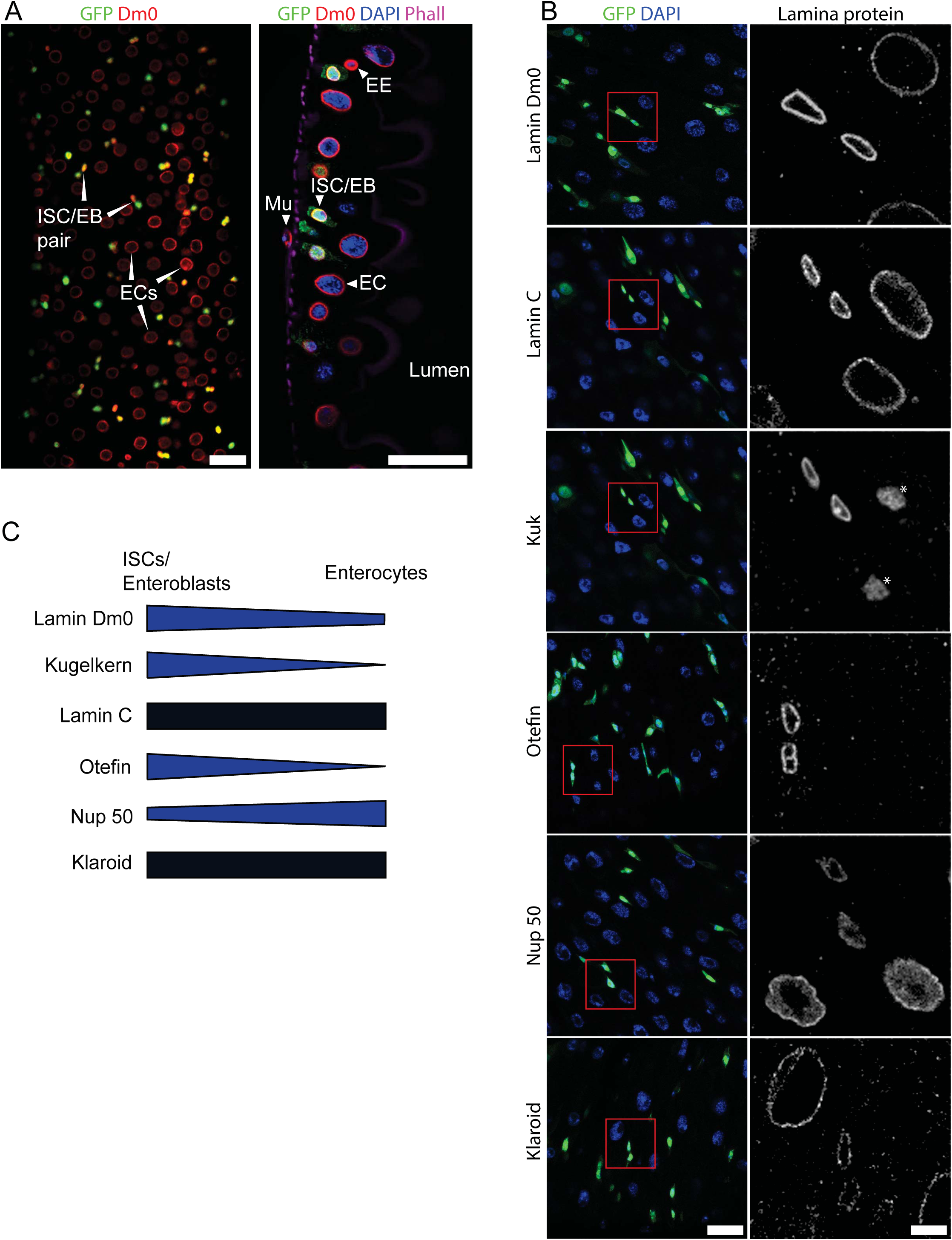
Dynamics of lamina proteins. (A) Fixed adult midguts stained as indicated for GFP, Lamin Dm0, DNA (DAPI) and F-actin (Phalloidin). Onview and sideview of the midgut. Immunostaining for Lamin Dm0 (red). Intestinal stem cells (ISC), enteroblast (EB) (green, GFP), polyploid nuclei of enterocytes (EC), Viceral muscle cell (Mu) and Enteroendocrine cell (EE). Scale bars: 25 μm (B) Fixed WT guts stained for lamina proteins (gray) inset, red box. ISC/EBs (green, GFP), DNA (blue, DAPI). Asterix: Unspecific staining of a nucleoplasmic structure by Kuk antibody. This staining is also observed in kuk deficient guts. Scale bar: 25 μm, 4x enlarged inset 5 m (C) Schematic of lamina protein dynamic in ISC/EBs versus ECs.

ISC proliferation is regulated by a set of signalling pathways that originate from adjacent cells. ECs and EBs promote ISC division by secretion of cytokines or mitogens which induce JAK/STAT, EGFR/Ras/MAPK, Hippo and Wg/Wnt signalling in the ISCs [2, 3, 6, 16, 34]. Viceral muscle cells induce canonical Wnt, JAK/STAT and EGFR signalling to promote stem cell maintenance over a longer time span and insulin signalling in response to food uptake [20, 21, 22, 39, 9]. ISCs and Enteroblasts often stay in close proximity and regulate their identity and proliferation by Delta/Notch signalling [23, 24, 25].

Here we investigate whether and how farnesylated proteins Lamin Dm0 and Kuk influence proliferation of ISC in adult *Drosophila*. We find that both Lamin Dm0 and Kuk strongly suppress ISC proliferation during homeostasis and regeneration. Lamin Dm0 antagonizes the regulation of the cell cycle in that Jak/Stat signalling is suppressed on the level of transcription as indicated by normalization of half of the Jak/Stat target genes in ISCs.

## 2 Results

As a starting point, we analysed the dynamics of lamina proteins in the midgut of adult *Drosophila* [35, 32, 31]. Generally, B type lamins are assumed to be ubiquitously expressed, whereas A-type lamins are assumed to show cell type specific expression [10]. We stained fixed guts with a panel of antibodies specific for lamina proteins to establish their dynamics during differentiation. ISC/EBs were marked by GFP, enterocytes recognized by cell size and polyploid nuclei. We found that Kuk and the LEM domain protein Otefin specifically marked ISC/EBs. Contrary to expectation we found that Lamin Dm0 showed stronger staining in ISC/EBs than EC, whereas the A type lamin, Lamin C was uniformly stained, as is the SUN domain protein Klaroid. The nuclear pore protein Nup50 showed a slight increase in nuclear staining of ECs compared to ISC/EBs (Fig. 1B, C).

Dm0 and Kuk inhibit ISC proliferation Expression of Lamin Dm0 and Kuk in adult tissue such as muscle or fat body induces changes in nuclear morphology, increased DNA damage, decreased heterochromatin and reduced lifespan [4]. Here, Lamin Dm0 and Kuk were expressed, additionally to endogenous levels, specifically in ISC/EBs and marked clones (flipout) using temperature sensitive GAL4/GAL80 system [16]. We assayed proliferation by the size of marked clones in female flies. The relative size of the clones indicate overall proliferation in the period between clone induction and analysis. The relative size of clones gradually increased over about 2 weeks, when the gut finally consisted of only clonal cells (Fig. 2A).

**Figure 2.**
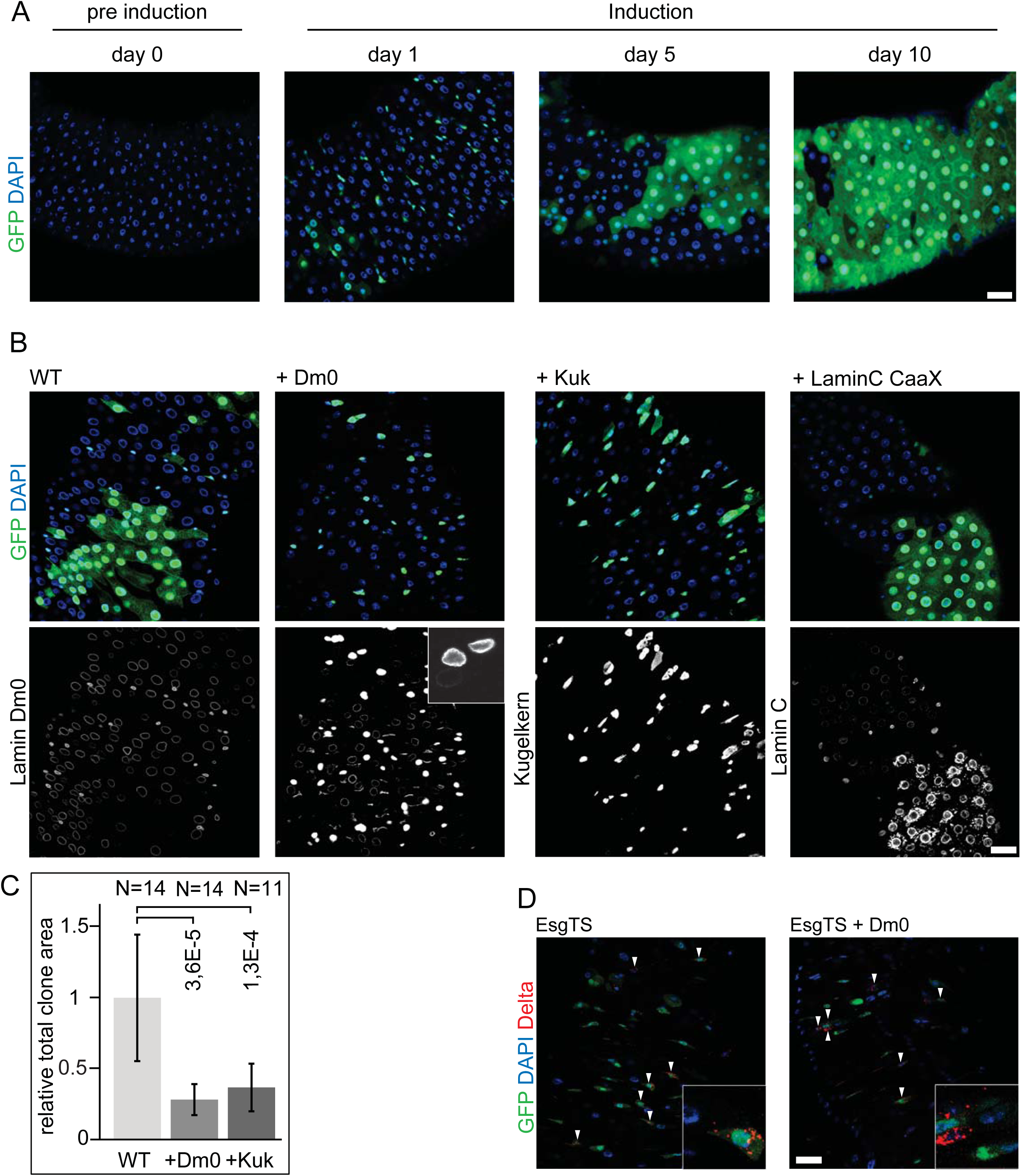
Lamin Dm0 inhibits ISC proliferation. Flipout clones, induced in ISC/EBs, (green, GFP) in adult female midguts fixed at indicated day after clone induction. The size of clones reflects the sum of proliferation during the induction time. (B) Midgut with flipout clones (green, GFP) fixed after 5 days of induction and stained for DNA (blue, DAPI) Lamin Dm0, Kuk and Lamin C (grey) as indicated. Clonal cells expressed GFP and in addition Lamin Dm0, Kuk or Lamin C CaaX as indicated. Immunostaining-image for Lamin Dm0 expression was oversaturated to make ECs visible and Lamin Dm0 levels comparable to WT. Inset for Lamin Dm0 3x enlarged without oversaturation. (C) Relative total clonal area after 5 days of induction expressing GFP (WT) Lamin Dm0 (+Dm0) or Kugelkern (+Kuk). Error bars: standard error. P value: Students T-test, two tailed, two-sample unequal variance. (D) Expression of GFP (EsgTS, 5 days) or Lamin Dm0 (EsgTS + Dm0, 15 days) in ISC/EBs (green, GFP, no flipout clones). Fixed midguts stained for DNA (blue, DAPI) and Delta (red). Arrowheads point to Delta and GFP positive cells. Insets 4x magnification. All scale bars: 25 μm.

Consistent with previous work [16], the turnover under standard laboratory conditions is about two weeks in females. In contrast, ISC proliferation was completely inhibited by expression of Lamin Dm0 and Kuk in ICS/EB and derived clones. The clones remained small, largely consisting of individual cells. Kuk was slightly less active than Lamin Dm0, as small clones formed after 10 days consisting of a few cells only. In both cases immunostaining showed increased levels of Lamin Dm0 or Kuk in clones (Fig. 2B and 2C), indicating effective overexpression. The phenotype is specific for Lamin Dm0 and Kuk, as expression of a farnesylated version of Lamin C did not affect formation of large clones and thus proliferation of ISCs.

Single cell clones with GFP (clonal expression) were still present after 15 days. These cells stained for Delta which suggests that they maintained their ISC identity (Fig. 2D). This indicates that long term exposure to high Lamin Dm0 levels does not alter ISC identity or induces loss of ISCs.

In addition to homeostatic conditions we also tested proliferation during regeneration of the gut after bacterial infection. Feeding flies with pathogenic bacteria such as *Ecc15*, induces a regeneration response, including a boost in ISC proliferation [7]. Following expression of Lamin Dm0 in ISC/EBs for 8 days and short starvation, flies were fed with *Ecc15* bacteria (Fig. 3A). We measured the proliferative response by the number of mitotic cells per gut (Fig. 3B). Compared to control flies, which were fed by *E. coli* (DH5α), infection by *Ecc15* bacteria significantly increased proliferation. Expression of Lamin Dm0 in ISC/EBs strongly reduced proliferation by about half (Fig. 3C). Our data show that expression of Lamin Dm0 in ISC/EBs suppresses their proliferative response and thus tissue turnover under homeostatic conditions as well as during regeneration following bacterial infection.

**Figure 3.**
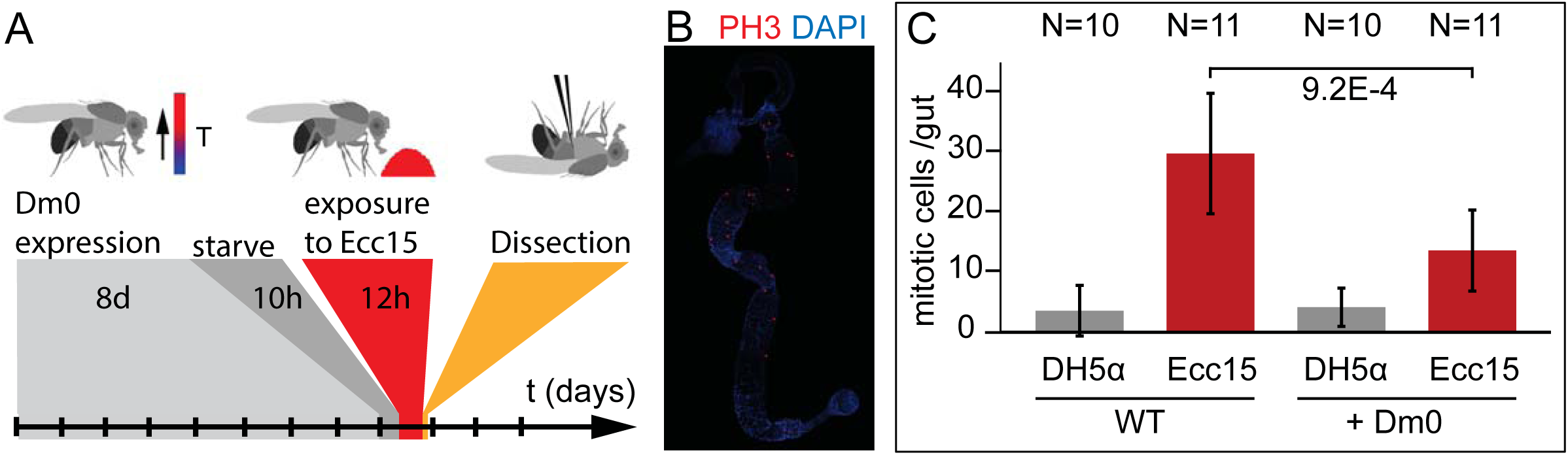
Lamin Dm0 inhibits *Ecc15* induced infection. (A) Experimental scheme for Lamin Dm0 expression and *Ecc15* infection. (B) Fixed gut stained for mitotic nuclei. (C) Mitotic cells per gut. Guts expressing GFP (WT) or Lamin Dm0 (+Dm0) infected with *E. coli* (DH5α) or pathogenic *Ecc15*. Error bars: standard error. P value: Students T-test, two tailed, two-sample unequal variance.

### Lamin Dm0 does not directly inhibit cell cycle progression

The activity of Lamin Dm0 may be due to specific modulation of the cell cycle, e.g. activation of a checkpoint and cell cycle stage-specific arrest. Alternatively, Lamin Dm0 may act indirectly by suppression of a proliferative signal. We employed the FUCCI system in a version adopted to *Drosophila* [40] (Fig. 4A and 4B) to score the distribution of cell cycle stages in ISC/EBs. A shift between stages would indicate a stage specific arrest. In standard guts we found about two thirds of ISC/EBs cells residing in G2 phase and one third in G1 phase. The proportion of cells in S phase was low, which is consistent with the low proliferation rate during homeostasis (Fig. 4C). Following *Ecc15* infection G1 and G2 phases are balanced and a small proportion of cells in S phase are observed. Expression of Lamin Dm0 did not change the distribution of cell cycle stages under homeostatic conditions and following *Ecc15* infection. Thus the anti-proliferative effect of Lamin Dm0 seems to be indirect, since no change in the distribution of cell cycle stages or stage-specific cell cycle arrest was detected.

**Figure 4.**
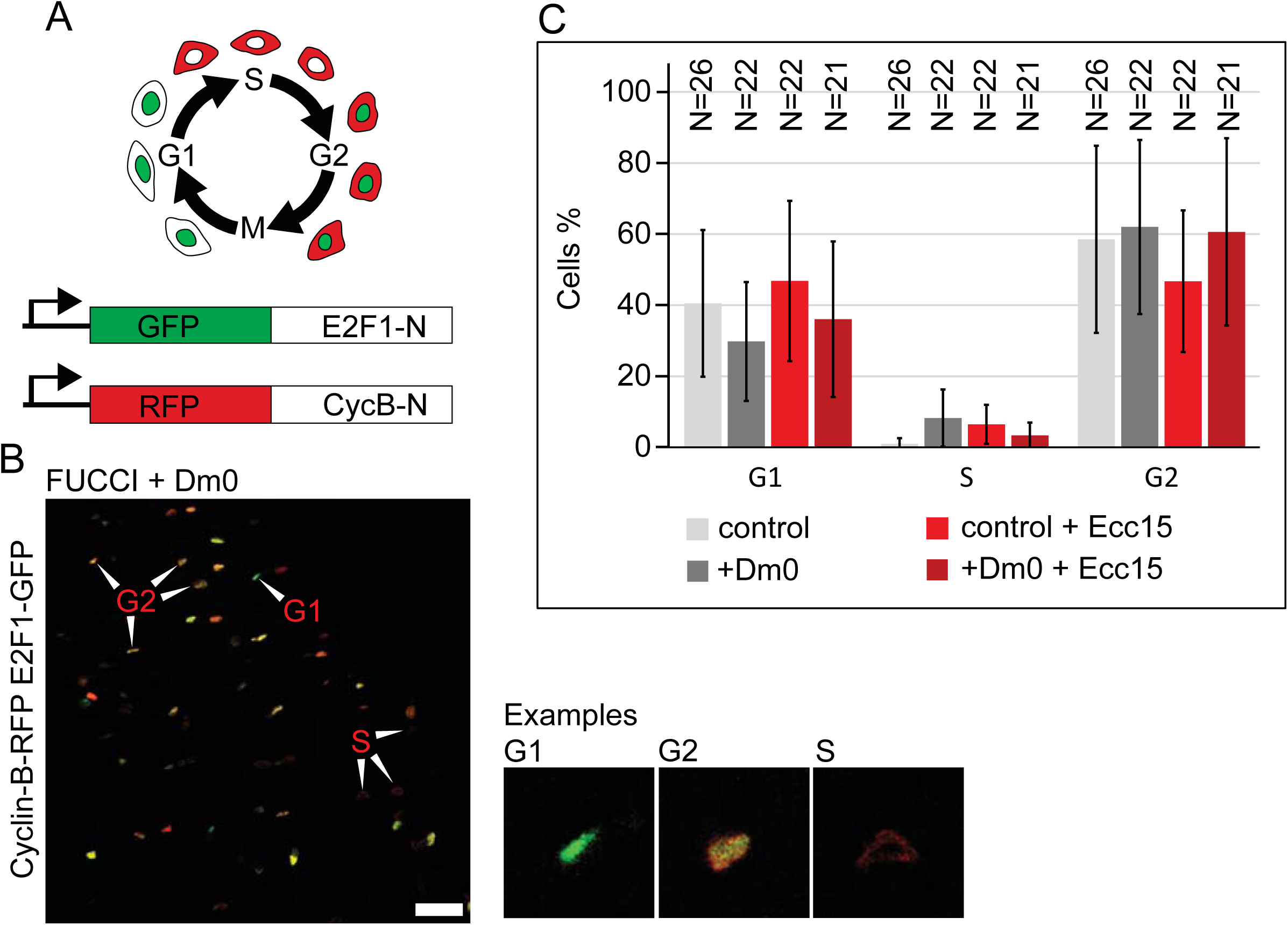
Lamin Dm0 does not change the cell cycle mode. (A) Schematic of the FUCCI system. Cell cycle with respective cells expressing GFP-E2F1-N or RFP-Cyclin B-N dependent on the cell cycle stage. GFP-E2F1-N/RFP-Cyclin B-N fusion proteins. (B) Example of FUCCI system in the midgut. Fixed gut with expression of GFP-E2F1-N (green), RFP-Cyclin B-N (red) or both (yellow) in ISC/EBs. Arrowheads point to exemplary cells for G1, G2 or S phase. Examples shown in high magnification. Scale bar: 25 μm. (C) Distribution of cell cycle modes. Wild type guts (control) and guts expressing Lamin Dm0 (+Dm0) and FUCCI reporters were infected with *Ecc15* (+*Ecc15*) as indicated. Error bars: standard error.

To further support a specific regulatory function on the cell cycle, we sought for a proliferative situation independent of Lamin Dm0. Activation of the Notch/Delta pathway is a potent inducer of proliferation. Depletion of Notch by RNAi induces a tumor-like proliferation of ISC cells [25]. We depleted Notch in ISCs/EB cells, which as expected led to strong ISC proliferation, as indicated by the clone size and high number of small cells. A similar phenotype was observed with concomitant expression of Lamin Dm0. Strong staining of Lamin Dm0 showed that expression of Lamin Dm0 was effective (Fig. 5). As Notch/Delta induced proliferation was observed while Lamin Dm0 was overexpressed, Lamin Dm0 appears not to directly target the cell cycle machinery but rather an upstream regulatory signal independent of Notch/Delta.

**Figure 5.**
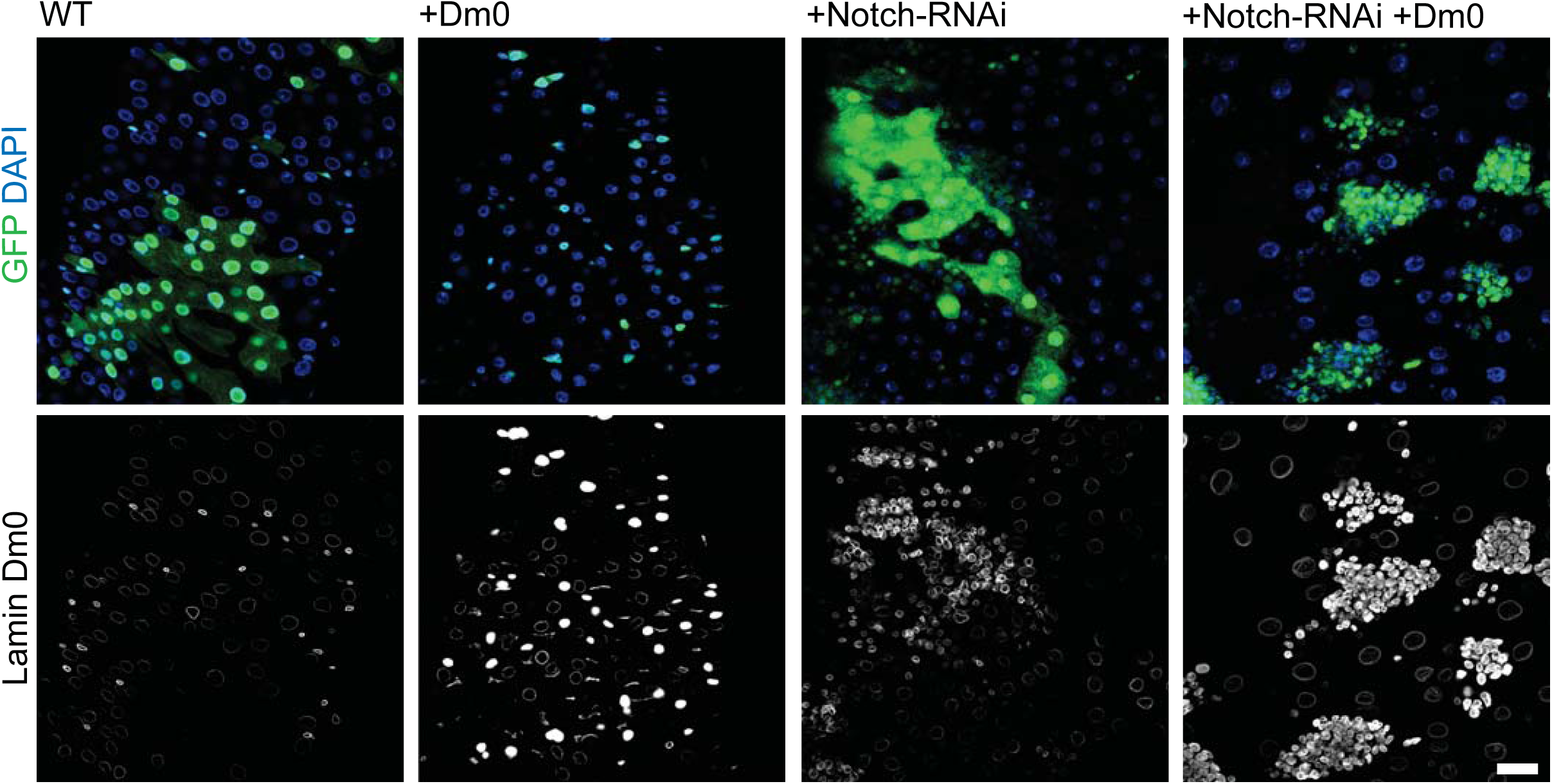
Lamin Dm0 does not inhibit Notch/Delta induced proliferation. Fixed guts with clonal expression of Lamin Dm0 and Notch RNAi as indicated. Flipout clones (green, GFP), DNA (blue, DAPI), Lamin Dm0 (grey). Scale bar: 25 μm.

### Lamin Dm0 and Kuk inhibit Jak/Stat induced proliferation

The Jak/Stat pathway is one of several pathways that transduce proliferative signals in the midgut. Jak/Stat signalling is induced by the cytokine Unpaired (Upd) and its receptor Domeless (Dome) and plays an essential role in controlling ISC proliferation and tissue turnover under homeostatic and regenerative conditions. We induced ISC proliferation by Upd expression in flipout clones and assessed proliferation by clone size. This led to complete tissue turnover in less than five days, which is consistent with previous reports [16]. In contrast, almost only single cell clones and a strong reduction in clone size were observed following coexpression of Lamin Dm0, suggesting an strong suppression of Jak/Stat signalling by Lamin Dm0 (Fig. 6A and 6B). A corresponding suppression of Jak/Stat signalling was observed with coexpression of Kuk (Fig. 6B and Fig. S1).

**Figure 6.**
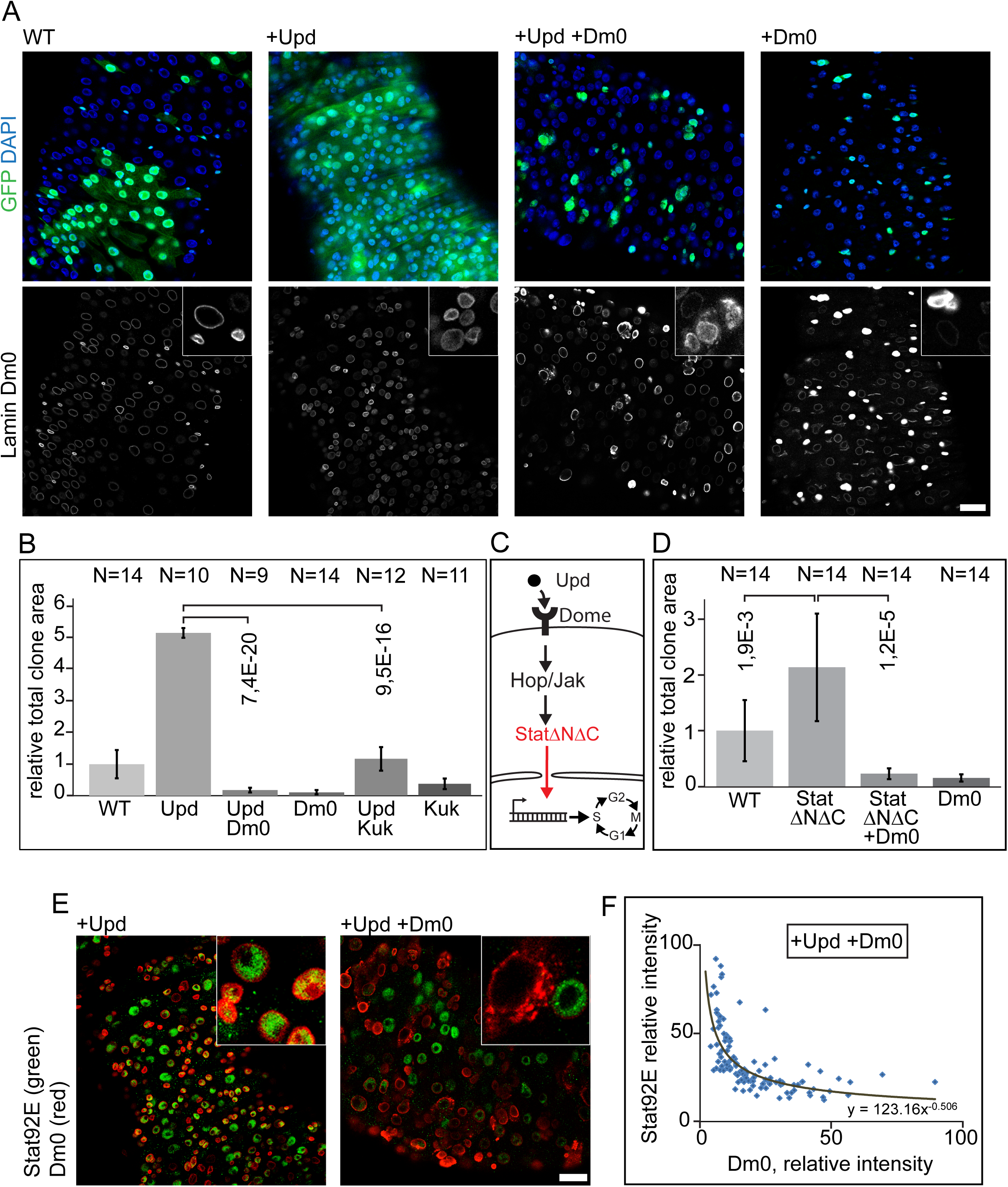
Lamin Dm0 and Kugelkern inhibit Unpaired induced proliferation. (A) Fixed guts with clonal expression of Unpaired (Upd) and Lamin Dm0 (+Dm0) as indicated. Flipout clones (green, GFP), DNA (blue, DAPI), Lamin Dm0 (grey). Inset, 4x magnification. 5 days of induction. (B, D) Relative total clonal area after 5 days of induction expressing GFP (WT) Lamin Dm0 (Dm0), Unpaired (Upd) or Kugelkern (Kuk). (C) Scheme of the Jak/Stat pathway with constitutive induction at the level of Stat by expression of StatΔNΔC. (E, F) Guts with flipout clones expressing Upd or Upd and Lamin Dm0 were fixed and stained for Stat (green) and Lamin Dm0 (red). Inset, 4x magnification. 5 days of induction. (F) Fluorescence intensities of Stat and Lamin Dm0 staining in clonal cells plotted against each other.All error bars: standard error. All P values: Students T-test, two tailed, two-sample unequal variance. All scale bars: 25 μm.

Lamin Dm0 may curb Jak/Stat signalling on multiple levels. It may have a direct effect on signalling by interfering with nuclear import of activated Stat, activation of Stat by Jak or impair the transcriptional activity of Stat. Alternatively, LaminDm0 may interfere indirectly by signal-independent effects on expression levels of Stat or Jak, for example. We tested whether Lamin Dm0 interferes with Jak/Stat signalling upstream or downstream of Stat in genetic terms. We expressed a constitutively activated form of Stat, StatΔNΔC [11] in ISC/EB flipout clones, which does not depend on upstream activation for its downstream function (Fig. 6C) and scored the effect on proliferation by relative total clonal area. We found a moderate but significant activation of proliferation by StatΔNΔC, which was suppressed by Lamin Dm0 coexpression (Fig. 6D). These data suggest that Lamin Dm0 interferes with Jak/Stat signalling on the same level or downstream of Stat.

Stat is generally regulated by phosphorylation, which leads to nuclear import [29]. We first stained midgut tissue with a Stat antibody recognizing both phosphorylated and unphosphorylated forms. We detected nuclear localization of Stat in ISC/EBs and overlapping staining with endogenous Lamin Dm0 following Upd expression (Fig. 6E). In contrast, nuclear localization and strong Stat staining was lost in midgut tissue with Lamin Dm0 expression in ISC/EBs. We detected an almost exclusive distribution of either Stat or Lamin Dm0 staining. Quantification of both staining intensities showed an inverse distribution with high Lamin Dm0 levels associated with low Stat levels and vice versa (Fig. 6F). We used an antibody specific for the active, phosphorylated form of Stat, pStat. As expected, pStat staining showed anti-correlation with Lamin Dm0 staining in Upd and Lamin Dm0 coexpressing flipout clones (Fig. S2). In control flies Stat and pStat positive cells occurred inside and outside of clones without prevalence. Upon clonal expression of Upd flipout clones cover nearly all midgut surface area and show strong Stat and pStat staining. Non clonal areas are either devoid of cells or might show cells with low or similar Stat levels as surrounding clones. Presumably due to non-autonomous activation of Jak/Stat. In Upd and Lamin Dm0 coexpressing flipout clones Stat and pStat staining is dominant outside of clones while being at least strongly reduced in clones (Fig. S2). These data suggest that Lamin Dm0 affects Stat protein levels in our experimental condition, which may be due to a transcriptional or post-transcriptional mechanism. A likely activity of Lamin Dm0 is on the level of transcription. To test this we established the transcriptional profiles of sorted ISC/EBs with and without Upd and Lamin Dm0 expression. Following isolation, midgut tissue with GFP labelled ISC/EBs cells was dissociated and sorted by FACS. Immediately after sorting RNA was isolated and subjected to next-generation sequencing (Fig. 7A).Transcripts were assigned to the annotated *Drosophila* genome and relative averaged expression levels were calculated. We found that expression of Upd in ISC/EBs lead to an at least twofold upregulation of 224 transcripts and at least twofold downregulation of 374 transcripts (Fig. 7B, Fig. S3A). Expression of Lamin Dm0 lead to the downregulation of 1027 transcripts and upregulation of 673 transcripts (Fig. S3A). To identify a specific effect of Lamin Dm0 on the target genes of Upd/Jak/Stat signalling, we asked which Jak/Stat target genes would be changed by coexpression of Lamin Dm0. Surprisingly about half of the Jak/Stat target genes were normalized to levels within the twofold threshold (Fig. 7C). Sorting of the Jak/Stat target genes according to the degree of up-or downregulation shows, that both strong and weak target genes were normalized by Lamin Dm0 (Fig. 7C). Only the very strong target genes as a class remained outside of the 2x threshold. In few cases Lamin Dm0 even reverted upregulated transcripts of considerably high copy numbers to a level of significant downregulation.

**Figure 7.**
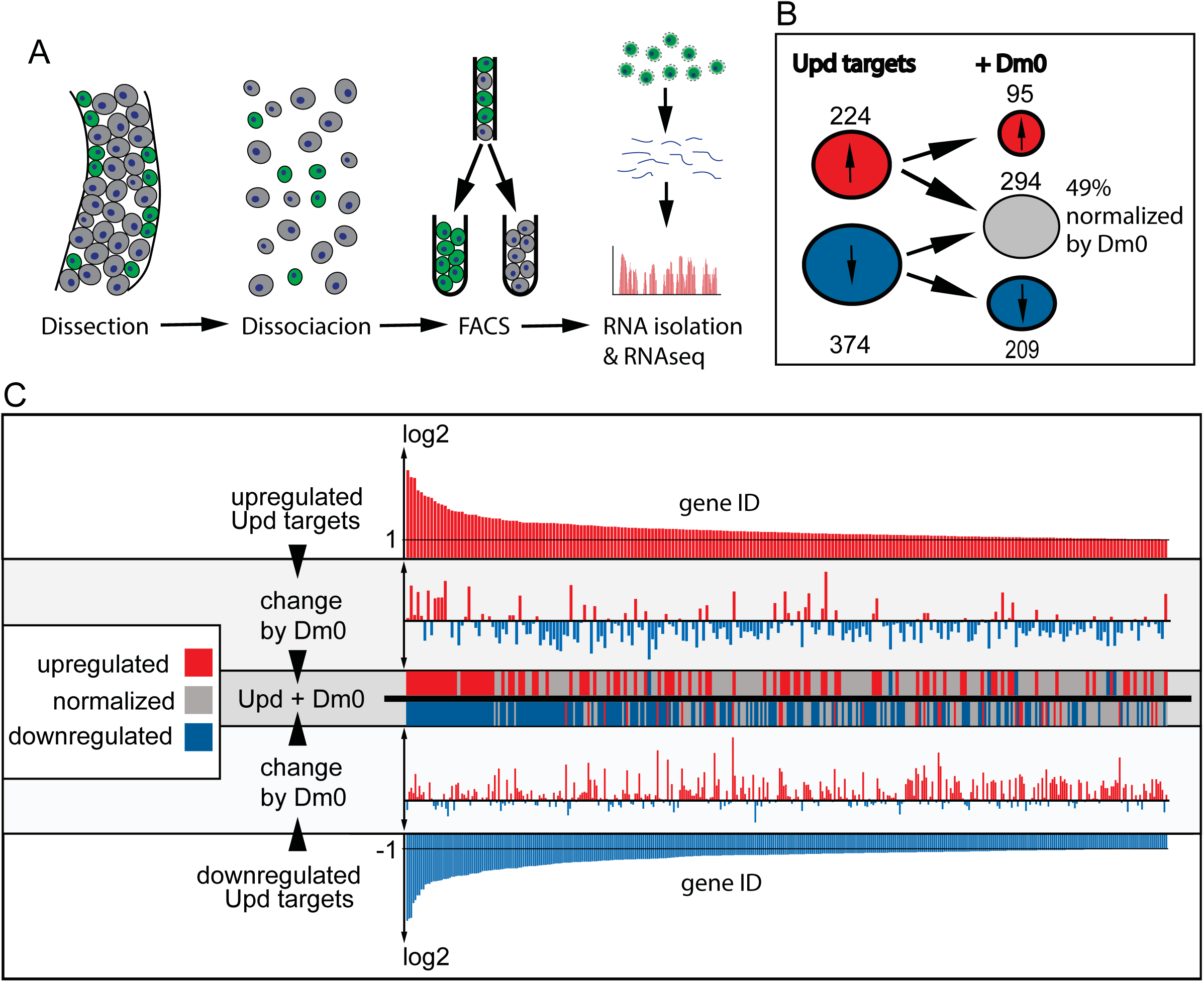
Lamin Dm0 normalizes expression of Jak/Stat target genes. (A) Experimental scheme. (B) Number of Upd induced up-(red) and downregulated (blue) transcripts with (+Dm0) or without Lamin Dm0 coexpression in ISC/EBs. (C) Change of Upd specific transcripts by Lamin Dm0 expression. Upper and lower panel show upregulated (red) and downregulated (blue) Upd target genes sorted by changes in expression level. Middle panels show changes of each target gene upon Lamin Dm0 coexpression. Red indicates upregulation (higher than 2 fold), blue downregulation (lower then 2 fold) and gray normalization.

These data suggest that Lamin Dm0 has a distinctive, target specific, effect on the transcription profile of Jak/Stat target genes. It is therefore unlikely that Lamin Dm0 affects Stat levels by reduced nuclear import because this would result in a general normalization of all Jak/Stat targets.

Among the Jak/Stat target genes whose expression was normalized by Lamin Dm0 coexpression was the Upd receptor Domeless (Dome). Dome is a Jak/Stat target with a 2.5-fold increased expression upon expression of Upd. Coexpression of Lamin Dm0 and Upd normalized Domeless expression to a factor of 1.2. (Fig: S3B). Our transcriptional data indicates that Lamin Dm0 might act as a normalizing, rather than outright inhibiting, force on Jak/Stat signalling. The normalized levels of Dome may contribute to the anti-proliferative effect of Lamin Dm0.

Lamin may normalize the transcription response to Jak/Stat signalling by recruiting parts of the genome to the nuclear periphery. It has been demonstrated previously that Lamin Dm0 interacts with specific regions of the chromosome. These so-called Lamina associated domains (LADs) are regions enriched in heterochromatin [37]. They are associated with gene silencing and potential regulation of developmental genes [27, 13]. To test an involvement of LADs in normalization of Jak/Stat targets by Lamin Dm0, we asked how many of the normalized target genes fall into LADs. Of genes that had significantly downregulated gene expression upon Dm0 expression 337 were found in LADs while 606 were found outside of LADs (Fig: S3C and S3D). So 35.7% of genes that had significantly reduced expression upon Dm0 expression were located in LADs. A similar ratio for genes in LADs was found for genes that had significantly upregulated gene expression upon Dm0 expression (35.1%). However of all genes annotated in *Drosophila* 39.5% are found in LADs.

Thus genes affected by Lamin Dm0 expression are equally likely to be associated with LADs than the general average of all genes. These comparisons indicate that genes inside or outside of LADs are equally likely to be affected by Lamin expression.

### Lamin Dm0 null clones do not show altered proliferation

The induced expression of Lamin Dm0 and Kuk in ISC/EBs reveals an anti-proliferative activity of these genes. To test whether these genes also have a function in homeostasis and regeneration of the midgut tissue, we analysed loss-of-function situations. Kuk-null homozygous flies are viable and fertile [5]. We did not observe any differences to wild type flies in number of ISC/EBs cells, indicating that Kuk is not required for ISC proliferation. In contrast to Kuk, Lamin Dm0 is partly required for viability. Only few Dm0-null mutant flies reach the adult stage [26]. To test for a function of Lamin Dm0 in ISC proliferation, we induced positively marked clones of Lamin Dm0 mutant cells within a wild type gut. As Lamin Dm0 clones were induced only in some ISCs, the clonal cells were in competition with wild type cells. Clonal cells contained no or only low levels of Lamin Dm0 persisting from the clonal mother cell, as demonstrated by immunostaining. Clone size did not differ noticeably between wild type and Lamin Dm0 clones even after 15 days. This indicates that Lamin Dm0 has no essential and unique function in homeostasis of the midgut possibly due to functional redundancy with other lamina proteins. Similarly, we could not detect a function of Lamin Dm0 in regeneration following bacterial infection.

Firstly, we did not observe a difference in clone size following *Pseudomonas entomophila* infection 5 weeks after clone induction (Fig. S4A and S4B). Secondly, the proliferation was not different in Lamin Dm0 clones and wild type guts following infection with *Ecc15* bacteria for 10h (Fig. S4C). These experiments indicate, that Lamin Dm0 is not required for proliferation of ISCs in the midgut during homeostasis and regeneration or that residual protein levels, originating from the ISC that formed the mutant clone, are sufficient to maintain its function.

## 3 Discussion

Expression of farnesylated lamina proteins can cause ageing associated phenotypes and progeroid diseases in humans and animal models. For instance expression of permanently farnesylated LaminA (Lamin50/Progerin) causes the Hutchinson Gilford Progeria syndrome (HGPS) which strongly reduces life span and causes several cellular and physiological effects reminiscent of ageing. How the cellular effects translate into the physiological effects is unclear but it is hypothesized that impairment of stem cells function might play a significant role. HGPS was modelled in mice where expression of Prelamin A or Progerin induces similar effects and stem cell numbers were found to be reduced [12, 33]. However how expression of farnesylated proteins might affect stem cell function is not well understood.

We investigated the activity of the farnesylated lamina proteins Lamin Dm0 and Kugelkern in *Drosophila* flies. We have previously shown that induced expression elicits ageing associated phenotypes on cellular and organismal levels, including shortened life span [4]. We now employed this experimental system to analyse their activity in the well characterized stem cells of the midgut. During differentiation, ISC/EBs show a stereotypic dynamic of lamina proteins, with Kugelkern and Lamin Dm0 staining high in ISC/EBs and low or absent in differentiated EC. Induced expression of Lamin Dm0 and Kuk in ISC/EBs and in clones strongly suppressed ISC proliferation. Despite this clear activity, Lamin Dm0 and Kuk are not required for ISC proliferation. This might be due to functional redundancy with Lamin C or other lamina proteins. Similar to mice [19] *Drosophila* flies homozygous for null mutations in the B-type lamin (Lamin Dm0) were able to reach the adult stage and survive in very low numbers [26]. We detected a suppression of ISC proliferation under two conditions. During homeostasis, proliferation is low with a few mitotic cells per gut and an overall tissue turnover of about two weeks. In contrast, proliferation is high during regeneration, which we induced by bacterial infection.

The proliferation block by Lamin Dm0 is not due to a remodelling of the cell cycle. Using the FUCCI system, we found that the distribution of the cell cycle phases was not changed much by Lamin Dm0 expression. The majority of ISCs remained in G2 phase. This finding suggests that Lamin Dm0 does not directly interfere with the basic cell cycle machinery. It is conceivable that Lamin Dm0 expression would downregulate expression of a cyclin or upregulation of a cell cycle inhibitor, which would lead to a cell cycle arrest. In such a case one might expect a shift of the cell cycle stages to a single stage. Consistent with a more regulatory influence of Lamin Dm0 on the cell cycle is our finding that Lamin Dm0 does not suppress Notch/Delta induced proliferation. Activation of Notch/Delta, e g. by Notch RNAi, leads to tumor-like proliferation. Such a proliferative response was also observed in guts, in which Lamin Dm0 was overexpressed. This finding shows that ISC proliferation is possible even when Lamin Dm0 is overexpressed. Our data are consistent with a model in which Lamin Dm0 acts on one or several specific pathways that regulate the cell cycle machinery.

Multiple signals are involved in control of ISC proliferation. Among these pathways we focused on the Jak/Stat pathway, induced by the cytokine Un-paired. This pathway is active under homeostatic and regenerative conditions. In contrast to Notch/Delta induced proliferation, cytokine induced proliferation was fully suppressed by Lamin Dm0 expression. Lamin Dm0 may suppress Jak/Stat signalling in two ways. Firstly, we observe a down-regulation of Stat protein in cells overexpressing Lamin Dm0. Co-staining of gut tissue showed a strong anti-correlative staining pattern. As we do not observe an effect of Lamin Dm0 on the expression of Stat RNA, the down-regulation of Stat protein may be due to reduced translation or increased degradation of Stat. Secondly, Lamin Dm0 modulates the transcriptional response of Jak/Stat signalling. Half of the Jak/Stat target genes in isolated ISC/EBs cells were normalized by concomitant Lamin Dm0 expression. Interestingly among the Jak/Stat target genes that are normalized by Lamin Dm0 is the cytokine receptor Domeless. This suggests that among other effects, the missing upregulation of the cytokine receptor leads to a weakened sensitivity of ISCs for cytokine signals. Our data further suggests that Lamin Dm0 affects Jak/Stat targets in a target specific manor which excludes a general effect on Stat import.

We propose a model where Lamin Dm0 expression inhibits ISC proliferation without depleting them. The effect of Lamin Dm0 expression is mediated in two ways. Lamin Dm0 reduces Stat protein levels posttranslationally but also normalizes the expression of 49% of Jak/Stat target genes in a target specific manor. Among the affected Jak/Stat targets is the Upd receptor Domeless. Reduction of Domeless leads to increased insensitivity of ISCs to Upd which further reduces proliferation. (Fig. 8).

**Figure 8.**
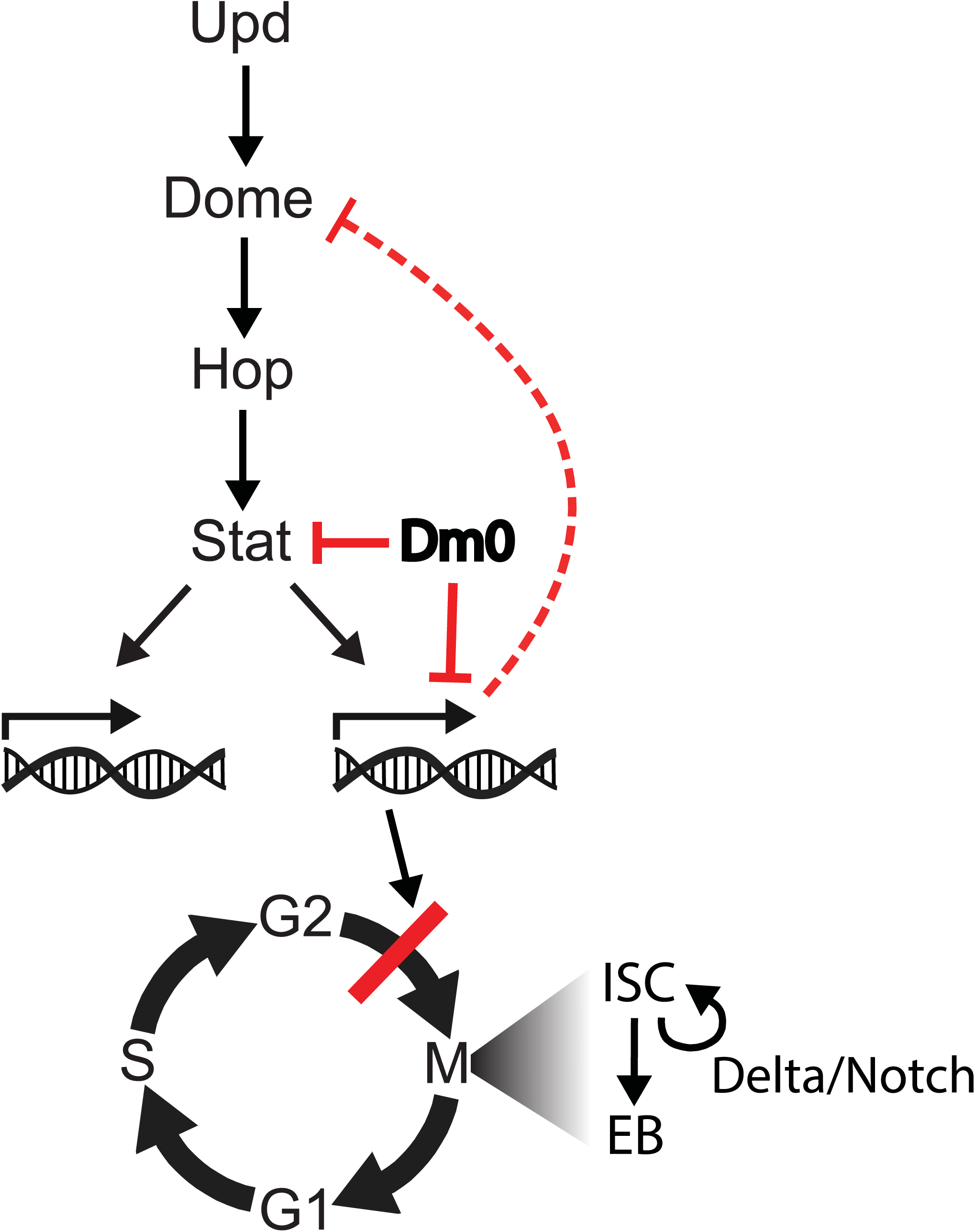
Model. Expression of Lamin Dm0 induces transcriptional changes that normalize Jak/Stat target genes. Normalization of Domeless expression may contribute to the suppression of Jak/Stat signalling. As a consequence of reduced Jak/Stat signalling, and possibly other pathways, proliferation is reduced. In addition to the transcriptional normalization Lamin Dm0 antagonizes Stat protein levels. Delta/Notch signalling control proliferation independent of Lamin Dm0.

## 4 Materials and methods

### 4.1 Genetics

#### 4.1.1 *Drosophila* stocks

In all experiments female *Drosophila melanogaster* fruit flies were used, aged 2 days to 5 weeks, dependent on the experiment.

**Table 1.** *Drosophila* stocks used in this work.

| Name | Genotype | Source | Ref. |
| --- | --- | --- | --- |
| EsgTS | Esg-Gal4, tub<br>GAL80TS, UAS-nlsGFP | Bruce Edgar | [23] |
| EsgTS; FO | Esg-Gal4, tub-Gal80TS,<br>UAS-nlsGFP;<br>act>CD2>Gal4, UAS-flippase | Bruce Edgar | [16] |
| Upd | UAS-Upd26.2 | Bruce Edgar | [16] |
| Lamin DmO | UAS-Lamin DmO |  | [14] |
| Upd;<br>Lamin | UAS-Upd26.2; UAS-Lamin DmO | this work |  |
| Lamin C CaaX | UAS-Lamin C CaaX | Georg Krohne |  |
| MARCM | hsFLP,UAS-CD8-GFP;<br>tub-Gal80FRT2L; tub-Gal4 | Bloomington<br>*Drosophila* Stock<br>Center |  |
| FRT2L | FRT2L | Bloomington<br>*Drosophila* Stock<br>Center |  |
| LamD395 | FRT2L::LamD395 | Yixian Zheng | [8] |
| FUCCI | UASp-GFP-E2F1,<br>UAS-mRFP1-NLS-CycB | Bloomington<br>*Drosophila* Stock<br>Center | [40] |
| Kuk | UAS-Kugelkern | this work |  |
| StatΔNΔC | UAS-Stat92ENC | Erika Bach | [11] |
| Notch RNAi | UAS-Notch RNAi | Vienna *Drosophila*<br>Resource Center |  |

#### 4.1.2 Flipout, MARCM clones

Flipout clones were induced, by temperature shift to 29°C, continuously for 5 days. MARCM clones were induced by temperature shift to 37°C, for 1 h, at 3 consecutive days. In both cases flies were collected 3-6 days after eclosion.

### 4.2 Histology

**Table 2.** Antibodies used in this work.

| Antibody | Source | Reference |
| --- | --- | --- |
| **Primary antibodies** |  |  |
| mouse--Lamin Dm0 | Harald Saumweb | [30] |
| mouse--Lamin C | Hybridoma Bank | AB528339 |
| rabbit--Kuk | Jörg Großhans | [5] |
| guinea-pig--Lamin Dm0 C-terminus | Georg Krohne |  |
| rabbit--Stat92E | Steven X. Hou | Hou et al., 1996 |
| mouse--pStat92E | Xinhua Lin | Zhang Y et al., 2013 |
| rabbit--PH3 | Millipore-Upstate | Cat. # 06-570<br>Lot # |
| mouse--PH3 | Millipore-Upstate | Cat. # 05-806<br>Lot # |
| Otefin | Georg Krohne |  |
| Klaroid | Michael Welte | Kracklauer MP et al. 2007 |
| Nup50 | Jörg Großhans | [5] |

| Secondary antibodies | Source |
| --- | --- |
| *Alexa-coupled antibodies used for* | Invitrogen |

**Staining and imageing:** Guts were fixed for 40 min in 0.2% Tween-20, 0.5% Nonidet P40, 8% Formaldehyde, in phosphate buffered saline (PBS). Guts were washed once quickly in PBS-T (PBS + 0.1% Tween) followed by permeabilization overnight in 0.5% Saponin, 0.5% Triton-X100, in PBS. Guts were washed once quickly in PBS-T and subsequently blocked 40 min with 5% BSA in PBS-T. The first antibody, in PBS-T, was added for 2 hours or overnight. Guts were washed 3x quickly in PBS-T followed by 1 hour wash in PBS-T. The secondary antibody was added for 2 hours or overnight. Guts were washed 3 times quickly in PBS-T followed by 1 hour wash in PBS-T. Guts were mounted on a glass slide in 47.5% Glycerin, 47.5% PBS, 5% 1,4-diazabicyclo[2.2.2]octane (DABCO), covered with a cover slip and sealed with nail polish. For quantification’s tile scan images of about 10 guts per genotype were made with an EC Plan-Neofluar 10x/NA03 objective. For all other pictures the LCI Plan-Neofluar 63x/NA1.3 objective was used.

**Confocal microscope: LSM 780 (Zeiss)**

### 4.3 Quantification of clone size

Relative clone size: Fiji Threshold function was used to define GFP positive clones, then the whole midgut area was encircled with the lasso tool and the mean fluorescence measured. Measurements of all genotypes were displayed relative to the average clonal area of the control flies.

### 4.4 RNA-seq

Four genotypes were processed: 1. EsgTS/CyO; Dr/TM3 (control) 2. EsgTS; Dm0/(Dr/TM3) (Dm0 expression) 3. EsgTS/Upd; +/(Dr/TM3) (Upd expression, induction of ISC proliferation) 4. EsgTS/Upd; Dm0/(Dr/TM3) (Upd expression together with Dm0 expression). From each phenotype100-150 guts were dissected and the intestinal cells dissociated by elastase treatment (4 mg/ml Elastase in dissociation buffer, both Sigma) and agitation. The dissociated cells were transported to the FACS facility and GFP positive cells sorted directly into RNA extraction buffer. The mix was transported back to the lab, the RNA extracted and frozen. Each genotype was processed 3 times in this manor (3x 100-150 guts) on different days. In the end three samples of extracted RNA per phenotype (12 samples) were given to the "Microarray Core Facility” for analysis.

100 to 150 guts, for each genotype, were dissected and transferred to a 1.5 ml Eppendorf tube containing 400 μl of PBS. 100 l of Elastase solution was added to reach a final concentration of 0.8 mg/ml. The guts were incubated for 1 hour at 27°C and agitated every 15 minutes, by pipetting up and down about 40 times. Afterwards the mixture was centrifuged for 10 min at 600 g at 4°C and the pellet resuspended in 500 μl PBS.

FACS was performed in the Scientific Flow Cytometry Facility of the University Medical Center Göttingen (UMG), affiliated with the Department of Hematology and Oncology. The dissociated gut cells were applied to a 50 m cell-sieve, removing enterocytes and cellular debris from the solution. The remaining cells were sorted for granularity, excluding damaged or clumped cells, and the presence of GFP, specific for ISCs and EBs. The cells were sorted directly into RNA extraction buffer (PicoPure RNA isolation kit). For the following procedure all reagents used were part of the Arcturus PicoPure RNA isolation kit.

After FACS, the sorted cells, in RNA extraction buffer, were incubated in a thermomixer at 42°C for 1 h. Then 500 l 70% RNAse free Ethanol was added, mixed well and the solution was applied to two extraction columns in volumes of 270 μl. The columns were centrifuged 2 min at 100 g, to bind the RNA, followed by 30 s at 16000 g. The flow-through was discarded, 100 μl Wash buffer 1 applied to each column followed by 1 min centrifugation at 8000 g.

The total RNA sequencing procedures were performed by the "Microarray Core Facility”. Medizinische Fakultät Georg-August-Universität Göttingen, using the Illumina “sequencing-by-synthesis” technology.

### 4.5 Bacterial infection

100mlLB-medium were inoculated with *Erwinia carotovora carotovora* (*Ecc15*), *Pseudomonas entomophila* (P.e.) or *E. coli* (*DH5α*) bacteria and incubated at 30C, overnight. The culture was centrifuged at 3466 g for 15 min, the pellet resuspended in 5 ml PBS + 5 % Sucrose and mixed with the upper food layer of a small fly food vial (without preservatives). A piece of filter paper was inserted into the vial and attached above the agar layer, without making contact, forming a platform for flies to rest and clean. To facilitate proper uptake of the solution the flies were kept without food and water for 6-10 hours, before being transferred to the prepared vial. The flies were kept exposed to the *Ecc15* for 12 h and then dissected.

## 5 Acknowlegements

B Edgar, SX Hou, X Lin, G Krohne, H Saumweber, H. Steffen, Y Zheng and the Developmental Studies Hybridoma bank created by NICHD of the NIH/USA and maintained by the University of Iowa, the Bloomington *Drosophila* stock center supported by NIH P40OD018537, the Genomic Resource center at Indiana University (supported by NIH 2P40OD010949-10A1) for initial experiments, discussions, materials or fly stocks. We thank the Transcriptome analysis laboratory of the Göttingen Medical School for RNAseq and transcriptome analysis. This work was in part supported by the Göttingen Center for Molecular Biology (equipment repair funds) and the Deutsche Forschungsgemeinschaft (GR1945/2-1 and equipment grant INST1525/16-1 FUGG).

## 7 Data availability

RNA seq data: GEO Submission number (GSE97307) [NCBI tracking system #18363487]

---

**Figure S1.**
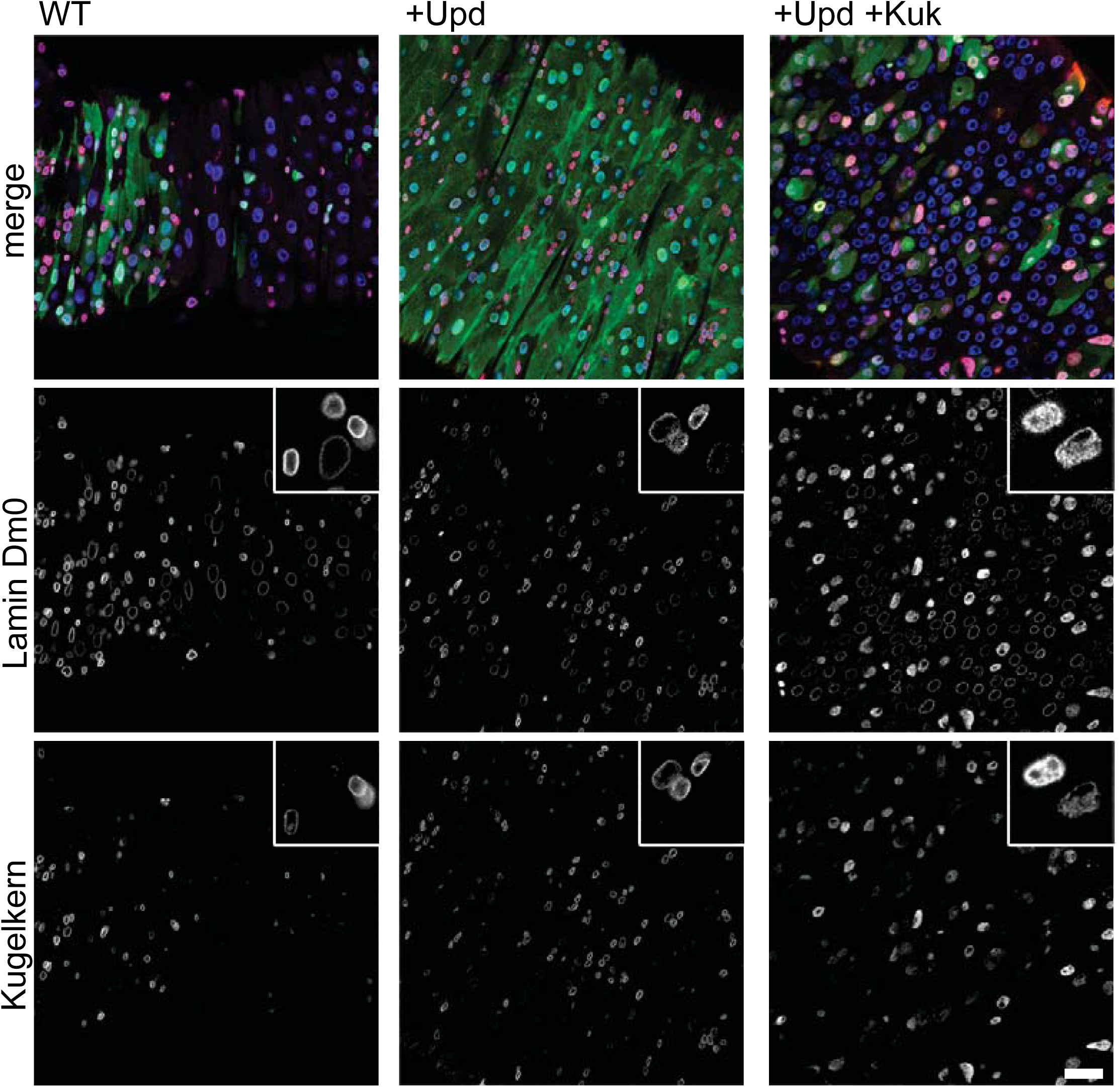
Kuk curbs Jak/Stat induced proliferation. Expression of GFP (WT) Unpaired (+Upd) and Kuk (+Kuk) in flipout clones as indicated. Flipout clones (green, GFP), DNA (blue, DAPI) immunostaining of Lamin Dm0 (purple, gray) and Kuk (gray). Insets show 3x enlarged examples of ISC/EBs. 5 days of induction. Scale bar: 25 μm.

**Figure S2.**
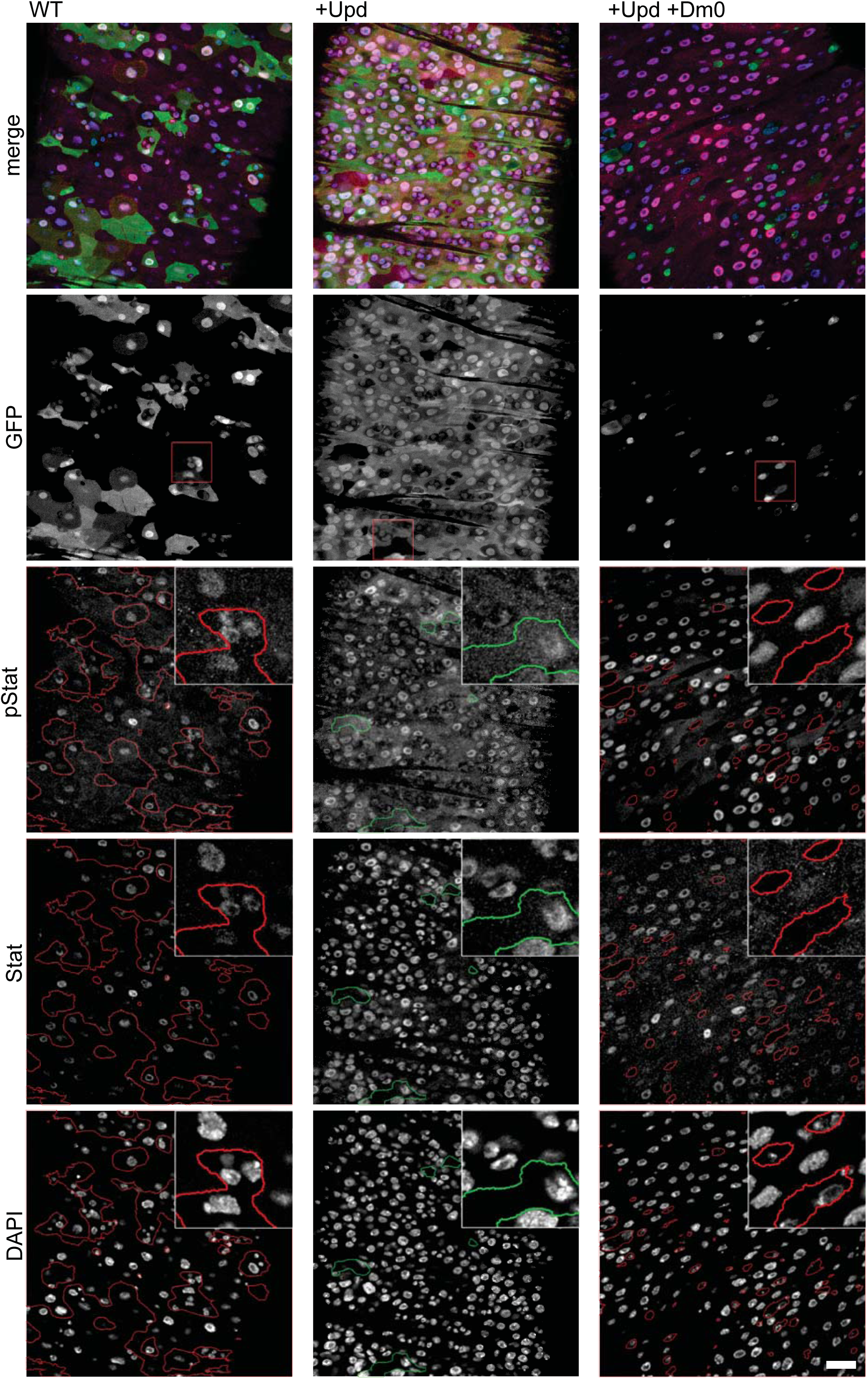
Lamin Dm0 antagonizes Stat and pStat protein levels. Expression of Upd or Upd and Lamin Dm0 coexpression in flipout clones, as indicated. Flipout clones (green, GFP), DNA (blue, DAPI) Stat immunostaining (purple). Single channels for GFP, Stat, DAPI (all in grey) as indicated. Clonal area is indicated in Stat and pStat channels by red outline, non-clonal area by green outline. Insets, 3x magnification of red boxed area in GFP channel. Scale bar: 25 μm.

**Figure S3.**
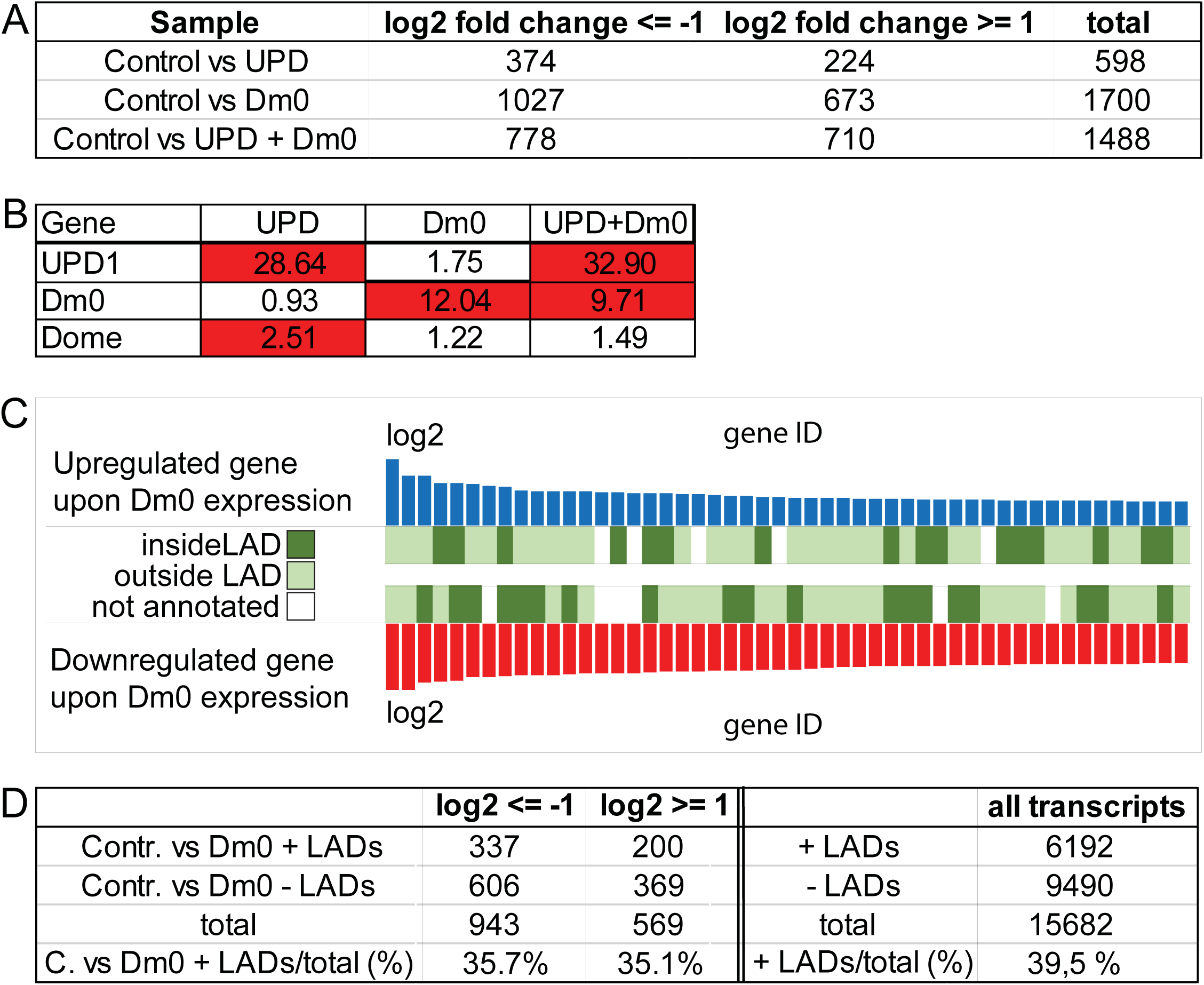
Transcriptional profiling. Results of RNAseq experiment of EsgTS (control) flies with expression of Upd, Lamin Dm0 or Upd and Lamin Dm0 in ISC/EBs. A: Number of transcripts that were downregulated or upregulated upon expression of Upd, Lamin Dm0 or both, in ISC/EBs with GFP expression, compared to only GFP expressing ISC/EBs. B: Selected genes that are upregulated (red) or unchanged/normalized (white). Numbers describe fold change in expression (not log2) C: Upregulated (red) or downregulated (blue) transcripts in relation to LADs (inside, outside or not annotated). Shown are the top 50 up/downregulated transcripts upon Lamin Dm0 expression. D: Left part of the table shows the number of transcripts that are downreg-ulated (log2 < =-1) or upregulated (log2 > =1) upon Lamin Dm0 expression and if the gene locus is inside a LAD (Contr. vs Dm0 + LADs) or outside a LAD (Contr. vs Dm0-LADs). The left part of the table shows all annotated *Drosophila* genes that lie in LADs (+LADs) and outside of LADs (-LADs) and the percentage of those that lie in LADs (+ LADs/total (%)).

**Figure S4.**
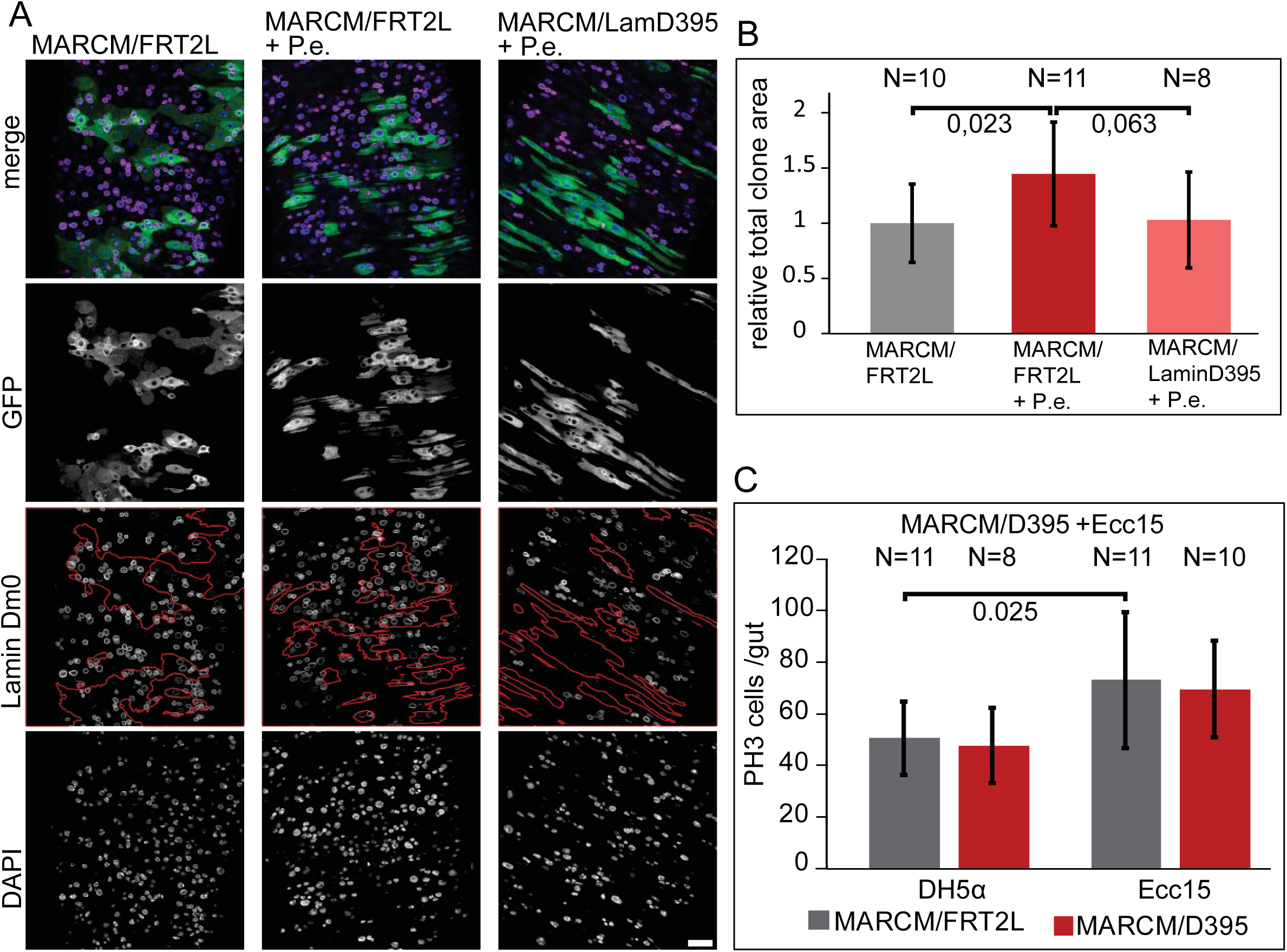
Lamin Dm0 is not essential for proliferation of ISCs. A, B: Guts with mitotic wildtype clones (MARCM/FRT2L) or loss of func-tion Lamin Dm0 clones (MARCM/LamD395) were infected with P.e. as indicated. Clones (green, GFP, gray), DNA (blue, DAPI, gray), Lamin Dm0 (purple, gray). Clones are marked by red outline in Lamin Dm0 staining. 5 weeks of clone growth was followed by P.e. infection as indicated. B: Relative clonal area. C: Mitotic cells per gut in guts with mitotic clones for wildtype or loss of function Lamin Dm0 clones. 4 weeks of clone growth followed by infection with DH5α or *Ecc15* as indicated. All error bars: standard error. All P values: Students T-test, two tailed, two-sample unequal variance. Scale bar: 25 μm.

